# Met58 and di-acidic motif located at C-terminal region of SARS-CoV-2 ORF6 plays a crucial role in its structural conformations

**DOI:** 10.1101/2023.08.14.553212

**Authors:** Prateek Kumar, Kumar Udit Saumya, Taniya Bhardwaj, Rajanish Giri

## Abstract

Despite being mostly neglected in structural biology, the C-terminal Regions (CTRs) are studied to be multifunctional in humans as well as in viruses. Their role in cellular processes such as trafficking, protein-protein interactions, and protein-lipid interactions are known due to their structural properties. In our previous findings on SARS-CoV-2 Spike and NSP1 proteins, the C-terminal regions (CTRs) are observed to be disordered and experimental evidence showed a gain of structure properties in different physiological environments. In this line, we have investigated the structural dynamics of CTR (residues 38-61) of SARS-CoV-2 ORF6 protein, disrupting bidirectional transport between the nucleus and cytoplasm. Like Spike and NSP1-CTR, the ORF6-CTR is also disordered in nature but possesses gain of structure properties in minimal physiological conditions. As per studies, the residue such as Methionine at 58^th^ position in ORF6 is critical for interaction with Rae1-Nup98. Therefore, along with M58, we have identified a few other mutations from the literature and performed extensive structure modelling and dynamics studies using computational simulations. The exciting revelations in CTR models provide evidence of its structural flexibility and possible capabilities to perform multifunctionality inside the host.

## Introduction

SARS-CoV-2, a beta-coronavirus, codes for several accessory proteins and is necessary for its pathogenesis along with structural and non-structural proteins. The accessory proteins do not contribute to its structure, however, they are critical for viral evasion of host immunity, formation of virus-induced membrane network, and hijacking metabolic pathways of host cell ^1^. ORF6 is one such protein, having been investigated for its potent role as an interferon antagonist and a nuclear import/export inhibitor. ORF6 is a seven kDa small transmembrane protein with a C-terminal cytoplasmic tail. It suppresses the translocation of STAT1 and IRF3 transcription factors into the nucleus by cytoplasmic accumulation of importin-α/β nuclear transport proteins ^2^. It also abolishes the export of host mRNA from the nucleus, causing the downregulation of various pathways ^3^. Further, it has been speculated to play a major function in the induction of membrane compartments for viral replication. Supporting this role, ORF6 has been shown to co-localize with the markers for endoplasmic reticulum, Golgi, endosome, and autophagosome vesicles ^2^.

Interestingly, ORF6 interacts with the components of nuclear pore complex, Rae1 and Nup98. Mediated by the C-terminal region of ORF6, this interaction abrogates the docking of cargo-import/export protein complex, inhibiting nuclear trafficking ^3–6^. Addetia et al., have thoroughly investigated this interaction via three different constructs of ORF6 with mutations in its CTR. All three constructs show abrogated binding with the Rae1-Nup98 complex ^5^. The methionine residue at 58^th^ position in the C-terminal tail of ORF6 is speculated to facilitate the key interaction with Rae1-Nup98 complex. ORF9b, another accessory protein of coronaviruses, interacts with the C-terminus of ORF6. Some short motifs like _49_YSEL_52_ and di-acidic (_53_DE_54_) motifs play key roles in interactions of ORF6 with other partner molecules ^1,2^. These reports show the importance of terminal regions of proteins which are frequently involved in interactions with other proteins.

Here, aiming to understand the conformational changes in C-terminal region (residues 38-61) of ORF6, we investigated this region in isolation. We studied this region in the presence of artificial environments to establish it as an intrinsically disordered protein region. Moreover, the structural properties of ORF6-CTR are also investigated in different physiological environments in vitro, such as organic solvents, liposomes, micelle-forming substances, and molecular crowders. Through computational modelling studies on ORF6-CTR, the structural dynamics is also inferred at hundreds of nanoseconds scale.

## Method and Materials

### Peptide and chemicals

The ORF6-CTR peptide with sequence _38_-KNLSKSLTENKYSQLDEEQPMEID-_61_ was commercially synthesized and obtained from Genscript LLC, USA. Chemicals like SDS, TMAO, Ethanol, PEG 8000, Ficoll, were procured from Sigma-Aldrich, USA. Liposomes were purchased from Avanti Polar lipids, USA. Other chemicals like TFE, HFIP were procured from HiMedia Co. Ltd., India.

### Liposome preparation and size measurement

The liposomes used in this study i.e., DOPS (1,2-dioleoyl-sn-glycero-3-phospho-Lserine), DOPC (dioleoyl-phosphatidyl-ethanolamine), and DOTAP (1,2-dioleoyl-3-trimethylammonium-propane) are procured from Avanti Polar Lipids (Alabaster, AL). These were prepared as per the previously reported protocols followed by the size estimation using Dynamic Light Scattering ^7,8^.

#### Circular Dichroism spectroscopy

CD spectroscopy was performed using 1 mm quartz cuvette on JASCO J1500 spectrometer using previously reported protocol ^9^. All measurements were taken in triplicates and the software provided average spectra. All spectra were smoothened by less than 10 points for analysis and representation purposes using the FFT filter in Origin Lab software.

#### Structure Model and Molecular Dynamics

In absence of full-length or partial structures of ORF6, the structures were modeled using AlphaFold plugin in ChimeraX ^10,11^. The best models were further processed for refinement and MD simulations using Gromacs. All parameters were calculated using Charmm36 forcefield using previously used protocols ^9^. Similar parameters were used for all mutant structure models of ORF6-CTR.

## Results

### ORF6 C-terminal shows disorderness in isolation

The C-terminal (residues 38-61) of ORF6 protein has been examined through web-based predictors for its disorderness. Three different sequence-based predictors viz. IUPred, PrDOS, and PONDR VSL2B have shown that this region has a much higher disorder propensity than the threshold value of 0.5 (**Figure 1a**). Previously, we have analyzed the full-length ORF6 protein’s sequence which has also demonstrated the C-terminal region to be disordered. To validate these predictions, we have recorded a Far UV CD spectrum of CTR in isolation where the negative ellipticity peak lies at ∼198 nm, a signature spectrum for unstructured proteins/regions (**Figure 1b**).

**Figure 1:**
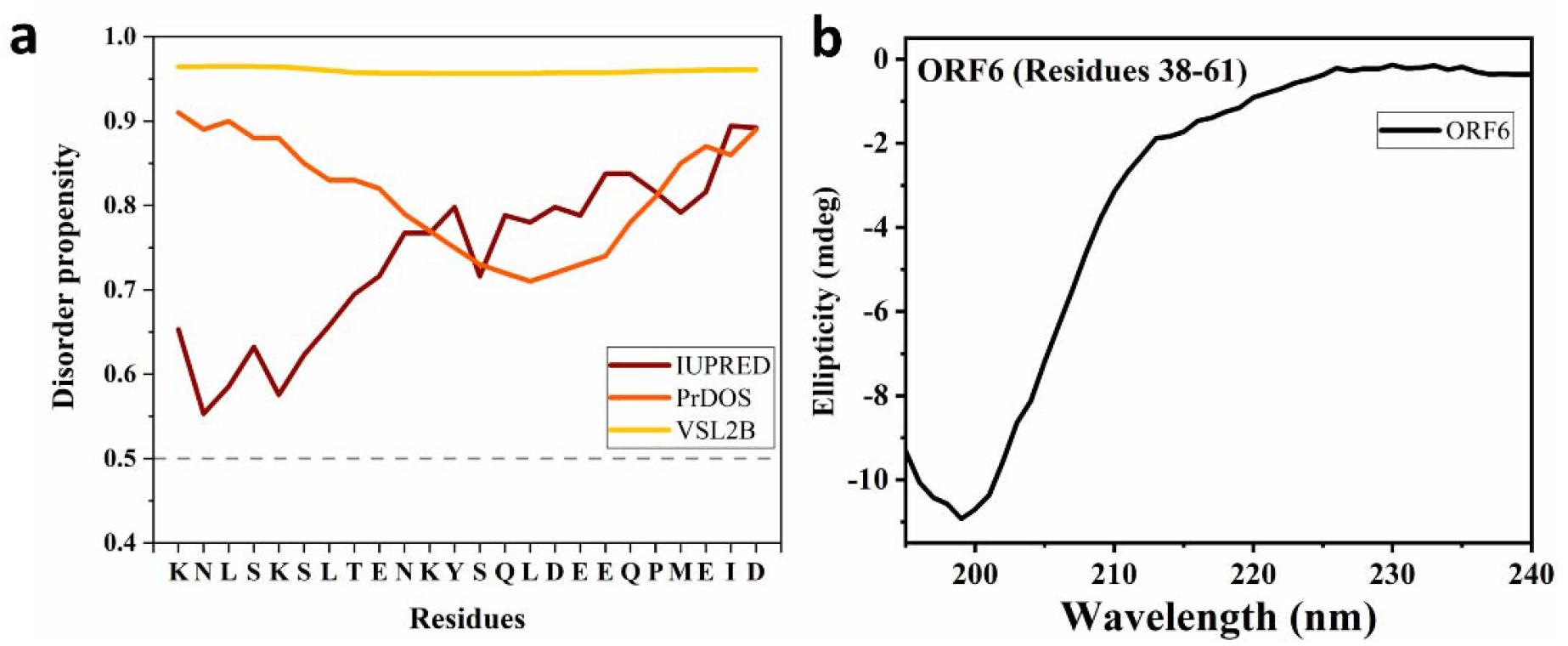
Disorder characteristics evaluation of ORF6-CTR.

### ORF6-CTR gains structure in organic solvents but not in molecular crowders

Organic solvents such as ethanol, TFE, and HFIP are used to investigate the effect of their surrounding environment on ORF6-CTR. There has been a gradual transition in secondary structure from disorder to order at the increasing concentrations of these organic solvents. In all graphic analyses, the helical structure is observed upon the addition of organic solvents with some other intermediate structures. Therefore, to deduce the precise changes from disorder (signature peak at ∼198 nm) to helical conformations (signature peaks at ∼208 and 222 nm), the 198 vs 222 nm graph is also plotted for all the measurements.

Specifically, more than 60% ethanol is required in the buffer composition to induce a proper helical structure of ORF6-CTR. In an absolute ethanol environment, the spectrum shift with two major peaks has been observed at wavelengths: ∼207 nm and ∼220 nm from the disorder spectrum at ∼198 nm (**Figure 2a**). A similar scenario can be interpreted from the 198 vs 222 nm plot, where structural transitions coincide at 70% ethanol (**Figure 2e**).

**Figure 2:**
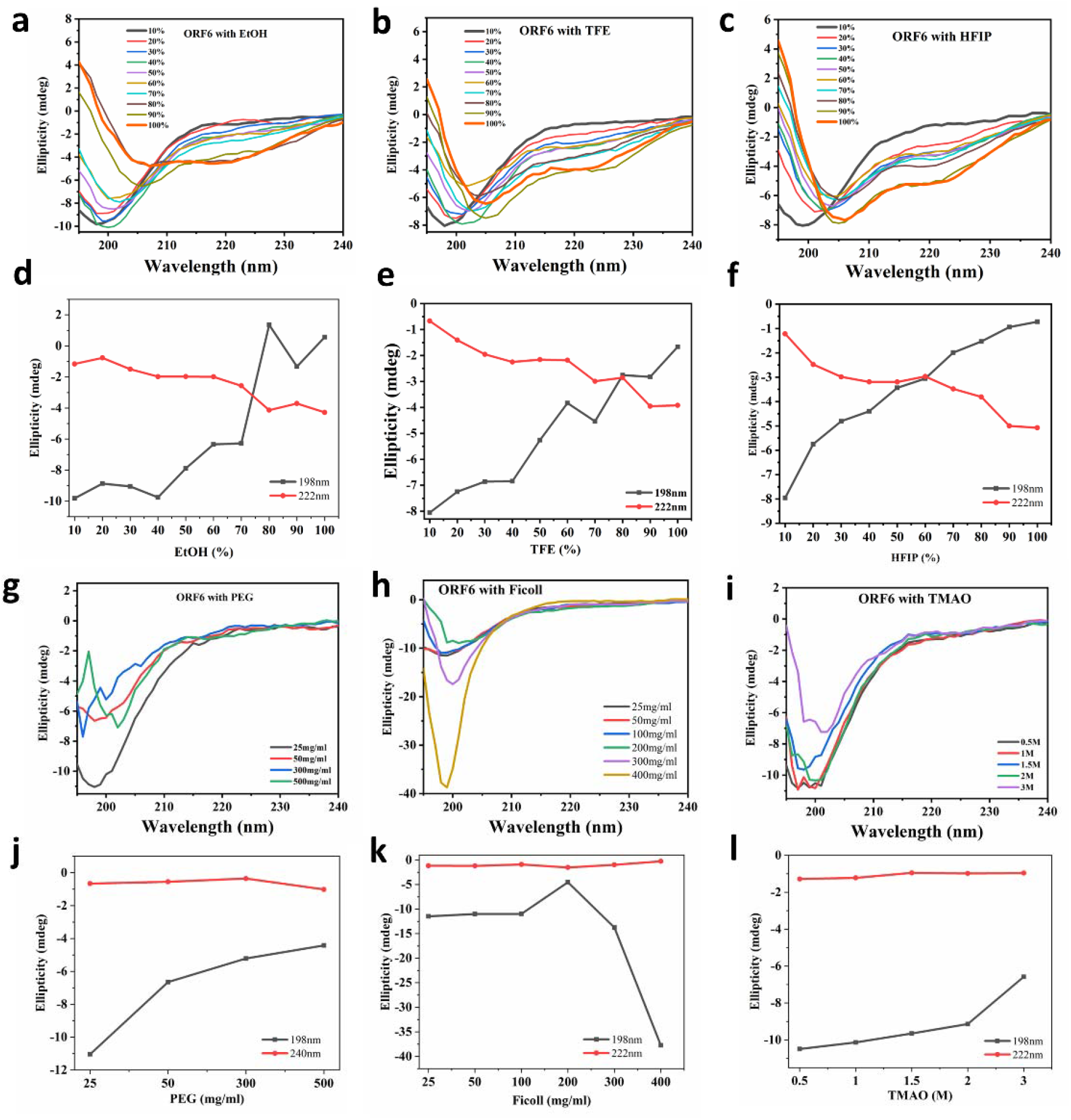
Using Far UV CD spectroscopy, the secondary structure evaluation of ORF6-CTR in the presence of organic solvents and molecular crowders.

Similarly, upon adding >80% TFE in the buffer, the peptide shows significant changes in its secondary structures. However, TFE is a helix inducer, but ORF6-CTR required >80% TFE in buffer to gain a structure. At 100%, the plot shows a typical helical structure spectrum with two negative ellipticity peaks at ∼206 nm and ∼222 nm (**Figure 2b**). These structural transitions in the presence of TFE are also evident from **figure 2f** where ellipticity at ∼198 nm is moving towards positive values while at ∼222 nm it shows the reverse order (**Figure 2f**).

In the presence of HFIP, the 24 amino acids long peptide has shown a gradual and firm transition from disorder to order starting at lower concentrations than the other two organic solvents. At only 20% HFIP, a dip in the spectrum is seen at ∼223 nm, which gets deeper with the increasing HFIP concentrations. Along with this dip, a gradual shift from ∼198 nm to ∼206 nm is also evident of structural transitions in the ORF6-CTR (**Figure 2c**). The 198 vs 222 nm plot also describes these conformational changes in **figure 2g**.

Further, the effect of molecular crowders such as PEG, Ficoll, and TMAO are also examined on the peptide using similar CD spectroscopy parameters. It is pretty evident that up to higher concentrations i.e., 500 mg/ml of PEG, 400 mg/ml of Ficoll, and 3M of TMAO, the ORF6-CTR does not gain any structure (**Figure 2g-2i**). Since no structural transitions are seen in the secondary structure, 198 vs 222 nm plots also do not signify any changes (**Figure 2j-2l**).

### Disorder persistency in lipid-mediated environments

ER membrane contains several lipids mediating multiple functions of newly translated proteins. Therefore, the presence of different liposomes (charged and neutral) in buffer conditions has been investigated on the peptide. Interestingly, neither positive/negative nor neutrally charged liposomes induce any structural changes in the peptide (**Figure 3a-3c**). ORF6-CTR remains disorder with a spectrum peak at ∼198 nm during most recorded spectra (**Figure 3e-3g**).

**Figure 3:**
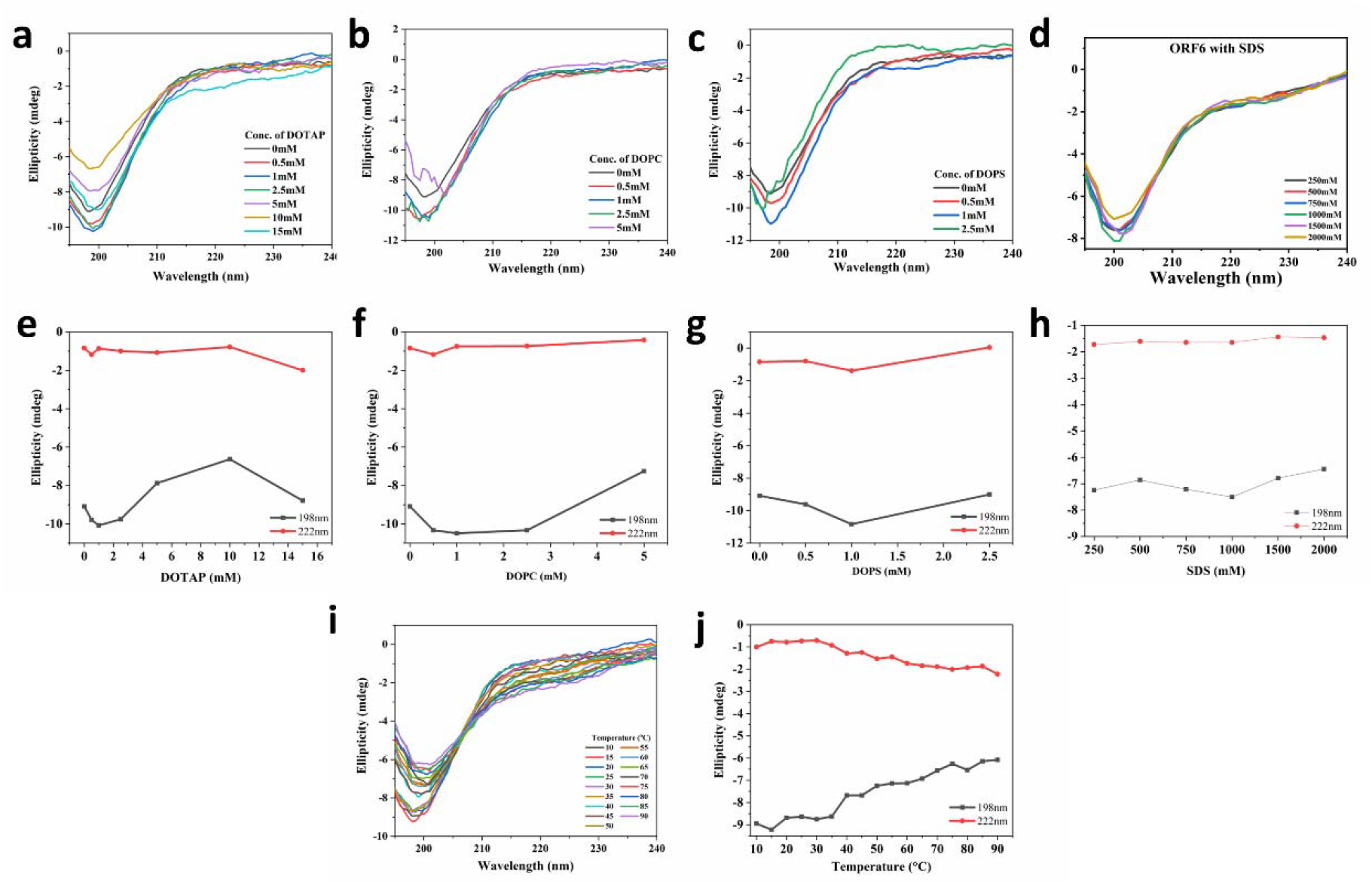
Secondary structure evaluation of ORF6-CTR in the presence of liposomes and SDS using Far UV CD spectroscopy. Mild contraction in peptide during increasing temperature in phosphate buffer.

Moreover, the micelle-forming environment by SDS could not affect the intrinsic properties of ORF6-CTR. At the concentration of 2M, the peptide remains disordered with no shifts in wavelength and no significant changes in the peaks at ∼198 nm (**Figure 3d & 3h**). However, in phosphate buffer and increasing temperature conditions, ORF6-CTR shows slight structural contraction with mild shifts and reduced peak intensities. These changes are consistent with increasing temperature and can be seen in **figure 3i & 3j**.

### Modelling of ORF6 C-terminal

So far, no full-length ORF6 or its C-terminal structure is available in the structure databases. However, a few structures with less than ten amino acids of C-terminal in complex with its host interacting partner Rae1-Nup98 complex are available. These structures show two small fractions of beta sheets and random coils. To overcome this, we have modeled the structure of the C-terminal region of ORF6 protein (**Figure 4**). The C-terminal structure possesses high disorder propensity, as per our sequence-based prediction and spectroscopy-based experimental outcomes. So, to look deeper into these outcomes, we modeled the structure using AlphaFold and found these outcomes to be in accordance with the above. The CTR structure contains a 3-residues helix in the middle and the rest is unstructured. All this together shows that the intrinsic structural properties of the C-terminal are disordered in nature.

**Figure 4:**
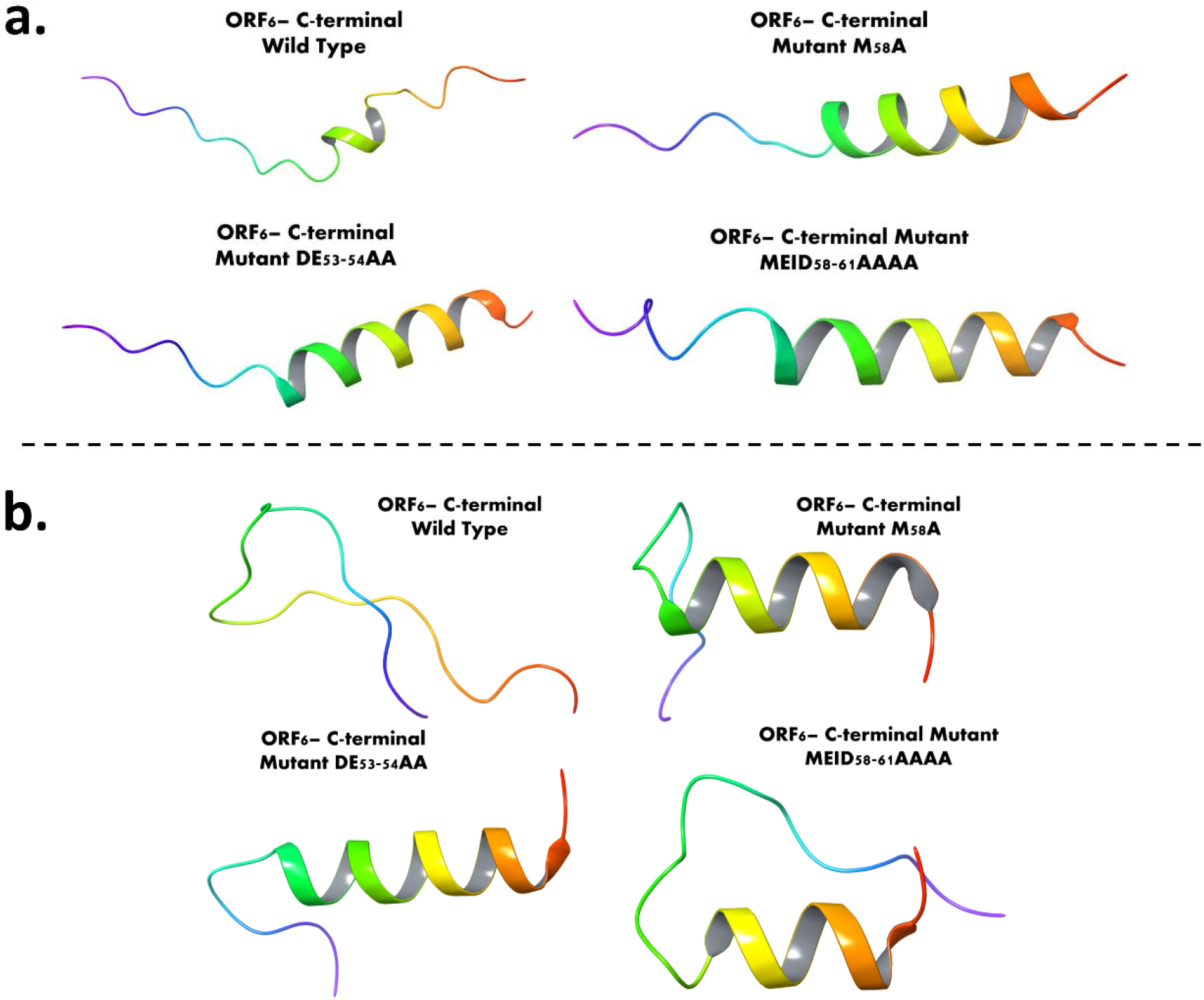
ORF6-CTR wild type and mutant type structures evaluation before and after MD simulations: **a**. AlphaFold based structure modelling of ORF6-CTR using wild type and mutated sequences (M58A, DE53-54AA, and MEID58-61AAAA). **b**. The last frames of 500 ns long wild and mutant type structure simulation trajectories.

Further, based on the literature, we have listed out some of the mutants in ORF6-CTR found to be functionally relevant and could affect the structure of ORF6. So, one such mutant is M58A, which has been studied to disrupt the interaction ORF6 with Rae1-Nup98. We modeled the structure by implementing this mutation in the CTR sequence and found an interesting phenomenon. **Figure 4** shows that with M58A mutation, the structure is modeled to be an α-helical with less unstructured region than the wild type. Another mutation occurs in its di-acidic motif at positions 53^rd^ and 54^th^ (_53_DE_54_). We mutated these residues to alanine at both positions and modeled the structure using AlphaFold. A similar phenomenon is observed with the M58A mutant. Lastly, we have modeled ORF6-CTR with another mutation in residues 58-61 (_58_MEID_61_ to _58_AAAA_61_). This, too, provides a helical structure with an unstructured tail region. So, this confirms that the di-acidic motif (_53_DE_54_) and M58 residues might be important for its functioning and be responsible for its disorder nature in native conditions.

### Met58 and di-acidic motif control the structural conformations of ORF6-CTR

Based on the literature, several mutations in ORF6-CTR were identified to be critical for its functions. Therefore, we performed MD simulations using Charmm36 forcefield in Gromacs to look deeper and its association with structure. For this purpose, we used the modelled wild-type and mutant structures from AlphaFold and prepared them in Chimera as described in the methods. A fascinating trend is revealed in simulations where the wild-type structure of ORF6-CTR has shown an entirely disordered-like structure which was previously containing a short helix, while the M58A mutant has shown a mainly ordered structure with *α*-helix of approximately 12 residues. The M58A mutant structure has maintained its structural integrity throughout the simulations, which is evident from trajectory analysis. However, the extended mutated region from 58-61 residues (_58_MEID_61_) has shown a loss of structure in comparison to modelled structure of the same region.

Similarly, di-acidic motif (DE) at positions 53^rd^ and 54^th^, when mutated and modelled through AlphaFold, the mostly *α*-helical structure was seen. Interestingly, after 500 ns long MD simulations, the structure has shown a complete structure integrity with less disorder composition compared to before simulation structure. As depicted in **figure 4**, the structure contains 15 residues long *α*-helix and then short disorder region. This confirms that di-acidic motif of ORF6-CTR is also responsible for structure-function activity of ORF6.

## Discussion

In structural biology, most of the proteins’ C-terminal regions (CTRs) are often excluded due to their missing electron density. Due to their structural flexibility, these CTRs are generally multifunctional in all life forms. These regions contain mostly short and few long motifs which allow them to function differently. Some examples of diverse functionalities are subcellular localization, protein-protein interaction through recognition of molecular features, protein-lipid interactions, etc ^12–15^. Also, the post-translational modifications such as glycosylphosphatidylinositol addition, farnesylation and geranylgeranylation at CTRs are found to be responsible for cellular processes ^16^. Interestingly, the C-terminus is also observed to be responsible for structural conformational transitions of MCAK protein ^17^. However, CTRs have not been the central focus of researchers across the globe still there are a substantial number of reports available on it. These regions are extensively studied in human proteins by identifying their roles in several cellular processes leading to critical diseases ^12,18,19^. Due to their diverse functions in cellular processes, the CTRs are also considered good therapeutic targets ^20,21^.

Apart from humans, the CTRs of viral proteins are also studied to some extent. As studied by Shukla et al., the CTR of Hepatitis A virus 2B protein is responsible for membrane interaction and possesses the pore forming capabilities in the membrane ^22^. An extensive mutations-based study by Jung et al. has revealed the importance of C-terminal of Core protein of Hepatitis B virus in RNA encapsidation and replication. The Core protein also possesses phosphorylation sites, which is confirmed by examining RNA encapsidation efficiency ^23^. In SARS-CoV-2, the nucleocapsid (N) protein is a highly disordered protein and its C-terminal possess dimer forming and nucleotide and protein interaction capabilities ^1,24^. Through Electron mobility shift assay, Zhou et al. confirmed that C-terminal domain of N protein can interact with single-stranded RNA and single/double-stranded DNA ^24^. Our previous reports on SARS-CoV-2 Spike and NSP1 proteins’ C-terminal domains also clearly demonstrate their role as molecular recognition elements and structural transition capabilities upon interacting with their binding partners ^7,9^. Possessing interaction motifs in the sequence allow them to interact with other viral and host proteins because of their intrinsic disorder nature, making them a potential therapeutic target ^21^.

Here we have observed that the ORF6-CTR is also intrinsically disordered in nature and cannot gain any structure upon interaction with any charged or uncharged lipids, molecular crowders. However, in the presence of high concentrations of organic solvents, it has shown the alpha-helical secondary structures. Modelling studies on CTR wild-type structures validate the experimental observations. Furthermore, the modelled structures with mutated sequences at positions _53_DE_54_, M58, and _58_MEID_61_ also contribute to the importance of CTRs in determining structural composition. The MD simulations results also portray similar outcomes where mutant-type structures hold more structural integrity than wild-type structures.

Altogether, we can speculate that ORF6-CTR’s disordered nature might be responsible for its interaction with ORF9b, and the residues such as _53_DE_54_ and M58 might be responsible for such interactions. Also, it is highly plausible that the interaction of ORF6 with Rae1 and Nup98 may depend on structural flexibility, as observed in the wild-type ORF6-CTR MD simulations. Indeed, these speculations need to be thoroughly tested in-vitro by mutating these residues and their interaction with ORF9b and other proteins. Moreover, such interactions might be targeted for the development of therapeutical strategies against SARS-CoV-2.

## Author Contributions

R.G.: Conception, design, and study supervision; P.K., K.U.S., and T.B.: acquisition of data, analysis, and writing of the manuscript.

## Acknowledgment

R.G. would like to thank the Department of Biotechnology grant (BT/11/IYBA/2018/06), MHRD-SPARC (SPARC/2018-2019/P37/SL), and Science and Engineering Research Board (SERB), India (Grant Number: CRG/2019/005603), and Indian Council of Medical Research (ICMR: 58/6/2020/PHA/BMS, and 52/04/2020/BIO/BMS). KUS and TB acknowledge ICMR and DST-INSPIRE for their fellowships, respectively.

## Conflict of interest

Authors declare no conflict of interest.

## Notes

### Competing Interest Statement

The authors have declared no competing interest.

